# Ultrashort treatment with telacebec alone and with companion drugs in immunocompetent and immunosuppressed mouse footpad models of Buruli ulcer

**DOI:** 10.1101/2021.06.22.449542

**Authors:** Oliver Komm, Deepak V. Almeida, Paul J. Converse, Till F. Omansen, Eric L. Nuermberger

## Abstract

The antimicrobial treatment of *Mycobacterium ulcerans* infection, or Buruli ulcer (BU), has a long duration and is therefore burdensome and linked to indirect costs for affected patients. The new antimycobacterial drug telacebec (Q203) has previously shown promising treatment-shortening potential in mouse models of BU. In the present study, we investigated the potential of Q203 to reduce the treatment duration further. The first experiment investigated the possibility of cure by one, three or five doses of Q203 (2 mg/kg) with or without a companion drug (bedaquiline, BDQ, clofazimine, CFZ, or clarithromycin, CLR) in immunocompetent BALB/c mice. The second experiment assessed the effect of five doses of Q203 with or without BDQ or CFZ on *Mycobacterium ulcerans* infection of immunocompromised SCID-beige mice with the aim to evaluate the contribution of host immunity to treatment efficacy. In BALB/c mice, a treatment duration as short as 3 days was sufficient to prevent relapse in nearly all footpads and a single dose of Q203 with or without BDQ or CFZ prevented relapse in approximately 50% of footpads. Unlike in BALB/c mice, a small percentage of SCID-beige mouse footpads were culture-positive after a treatment duration of five days, highlighting an important role of host immunity for *M. ulcerans* clearance. Our results confirm the marked potency and prolonged bactericidal and sterilizing effects of Q203, even in immunocompromised SCID-beige mice.

## INTRODUCTION

The neglected tropical disease Buruli ulcer (BU), which is caused by *Mycobacterium ulcerans* and which primarily causes skin and soft tissue lesions, is treated following WHO guidelines by a combination of 10 mg/kg rifampin (RIF, R) and 15-30 mg/kg clarithromycin (CLR, C) daily for 8 weeks, a long and burdensome treatment for the patients (1-5). This newer regimen, compared to the earlier regimen of RIF in combination with 15 mg/kg streptomycin (STR) daily for 8 weeks, avoids the risk of oto- and nephrotoxicity as well as the need for daily injections for 56 days. Nonetheless, the RIF+CLR treatment has disadvantages, such as frequent gastrointestinal intolerance, reduced CLR exposure due to a drug-drug interaction with RIF (6-9), and no reduction in the treatment duration of 8 weeks.

Accordingly, even though RIF+CLR is a clinically proven effective therapy for BU (4, 5), the search for a shorter oral treatment for BU is a priority for improved control of BU. Previous studies have sought to improve the efficacy of oral BU treatment. Experiments in mice have tried replacing CLR and/or STR with drugs like clofazimine (CFZ) or oxazolidinones or to give higher doses of rifamycins. Some of these new regimens allowed a reduction of the treatment duration, but no regimen was able to cure mouse footpad infections in less than 4 weeks (10-14).

The oxidative phosphorylation pathway was recently identified as a new drug target in *Mycobacterium tuberculosis* (15). Bedaquiline (BDQ) and telacebec (Q203, Q), two drugs acting at different steps on this pathway, proved to be efficacious for the treatment of BU (16-19). BDQ, an ATP synthase inhibitor, showed treatment shortening effects in regimens for drug-susceptible and multidrug-resistant tuberculosis (TB) (20-22). The recently rediscovered tuberculosis drug CFZ acts by being reduced by NDH-2, then reducing O_2_ resulting in the production of reactive oxygen species (23). Q203 is a mycobacterial cytochrome bc_1_:aa_3_ complex inhibitor that showed efficacy in mouse TB models and in an initial Phase 2 study in TB patients (24, 25). An important advantage of Q203 for the treatment of *M. ulcerans* is the lack of the alternative terminal cytochrome bd oxidase in this species (26), as a result of a nonsense mutation in the *cydA* gene (base substitution - 692G>A) (18). Q203 proved to be extremely potent against *M. ulcerans in vitro* with an MIC between 0.075 and 0.15 ng/ml (16, 18) and also as monotherapy in the mouse footpad infection model (17) as well as in 3- or 4-drug combination therapy with rifapentine, CFZ and/or BDQ (16). In both studies no CFU were detected 2 weeks after the end of a 2-week treatment. The addition of RIF at 10 or 20 mg/kg did not add a benefit (17).

In the present study using BALB/c mice we investigated if shorter treatment durations (1, 3 or 5 days) of Q203 at 2 mg/kg in monotherapy or 2-drug combination therapy with BDQ, CFZ or CLR, render mouse footpads culture negative, how long it takes to render the footpads negative, and if mice relapse after a 15-week observation period.

In the present study using immunocompromised SCID-beige mice we addressed the question of whether the continued reduction in CFU counts after completion of short courses of Q203 treatment in immunocompetent mice (16, 17) is primarily due to a prolonged antimicrobial effect of Q203 or due to the adaptive host immune response. Additionally, we tested the SCID-beige mouse footpads for emergence of bacterial resistance against Q203 and compared the results between groups, to assess the necessity of a companion drug for Q203 to prevent resistance.

## RESULTS

### Experiment 1: To determine if addition of a companion drug increases the sterilizing activity of Q203 and to find the shortest possible treatment duration

In this experiment, BALB/c mice were treated with 2 mg/kg Q203 (Q_2_) alone or in combination with one of the following drugs: 25 mg/kg BDQ (Q_2_BDQ_25_), 12.5 mg/kg CFZ (Q_2_CFZ_12.5_), or 100 mg/kg CLR (Q_2_CLR_100_). Drugs were administered once daily via oral gavage and each regimen was given for 1, 3 and 5 days. Cohorts of mice were followed for up to 15 weeks after the start of treatment (Table S1).

#### Footpad swelling and CFU counts up to 5 weeks post-treatment

On the day after infection, the mean (± SD) CFU count was 4.95 ± 0.38 log_10_ CFU/footpad. At the start of treatment (D0), 34 days after infection, the mean CFU count was 6.62 ± 0.34 log_10_ CFU/footpad, and the median swelling grade was 2 on a scale of 0-4 (27). Untreated mice subsequently exhibited increased footpad swelling and required euthanasia during the experiment (Fig. 1A), with the exception of one untreated mouse that survived until the end of the experiment without reaching the endpoint for euthanasia despite harboring 6.17 ± 0.86 log_10_ CFU/footpad at the end of the experiment. In mice treated with R_10_CLR_100_ for 10 consecutive days, the swelling decreased slightly to an average of 1.7 after one week of treatment and to 1.25 after 2 weeks. The decrease in swelling continued up to Week 4 and was maintained until Week 8, when a gradual increase in swelling started and ultimately resulted in four footpads having a swelling ≥3 by Week 14. The mean CFU count in R_10_CLR_100_-treated mice decreased to 3.81 ± 1.35 log_10_ CFU/footpad by Week 2 (Fig. 1B) but increased after the end of treatment to 5.49 ± 1.21 log_10_ CFU/footpad by Week 15.

**Fig 1.**
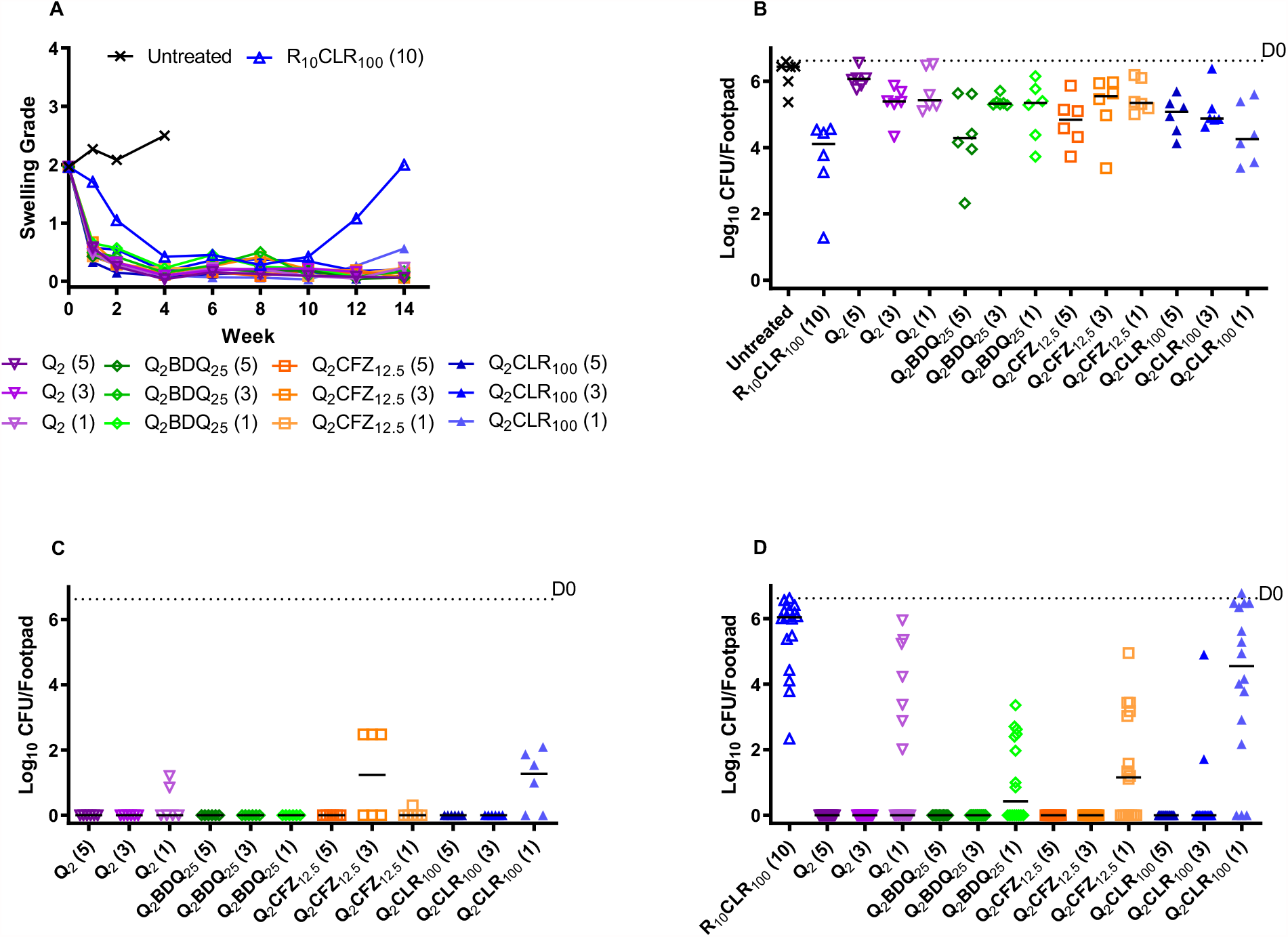
Footpad swelling grade and microbiological outcome in BALB/c mice in response to treatment. Median swelling grade in BALB/c mice during and after treatment with the indicated regimens and number of doses in parentheses (A). Log_10_ CFU/footpad one week after the start of treatment (B). Log_10_ CFU/footpad six weeks after the start of treatment (C). Log_10_ CFU/footpad 15 weeks after the start of treatment (D). R, rifampin, CLR, clarithromycin, Q, telacebec, BDQ, bedaquiline, CFZ, clofazimine. Numbers in subscript indicate the dose in mg/kg. D0, day 0 or the beginning of treatment.

As previously observed (17), Q203-containing regimens rapidly decreased footpad swelling, resulting in >98% of these mice having a swelling grade ≤1 by Week 1, significantly lower than the R_10_CLR_100_ control group (p<0.001). Among mice receiving Q203, 36% and 67% of the footpads showed no swelling by Weeks 2 and 4, respectively, with no significant difference between Q203-treated groups. In the first week, the superior effect of Q203 was not yet reflected in the footpad CFU counts. Five weeks after the end of treatment, all Q203-containing regimens given for 5 days resulted in culture-negative footpads (Fig. 1C). With the exception of three of 6 footpads from mice treated for 3 days with Q_2_CFZ_12.5_, footpads treated with a Q203-containing regimen for three days were all culture-negative. In the groups treated for only one day, 1 mouse (both footpads) that received Q203 alone had positive culture results, all mice receiving Q_2_BDQ_25_ for one day were culture-negative, one mouse receiving Q_2_CFZ_12.5_ for one day had a culture-positive footpad, and in the group receiving Q_2_CLR_100_ for one day, two mice had both footpads culture-positive at this time point. At Week 6, the CFU counts were significantly higher in mice treated with one dose of Q_2_CLR_100_ compared to one dose of Q_2_BDQ_25_ (p<0.03).

#### Relapse

Eight BALB/c mice of each treatment group were held for 15 weeks to evaluate relapse. Apart from the relapses observed in all R_10_CLR_100_-treated mice, we also observed relapse in all but 3 of the 16 footpads treated with a single dose of Q_2_CLR_100_. A single dose of Q203 resulted in 9 out of 16 footpads being culture-negative, a single dose of Q203 and BDQ resulted in half (8/16) of the footpads being culture-negative, one dose of Q_2_CFZ_12.5_ resulted in 7 of 16 footpads being culture-negative. At Week 15, the CFU counts were significantly higher in mice treated with one dose of Q_2_CLR_100_ compared to one dose of Q203 alone, Q_2_BDQ_25_ or Q_2_CFZ_12.5_ (p<0.02). When given for 3 days, Q203 with or without a companion drug prevented relapse in all footpads with the exception of 2 out of 16 footpads from mice treated with Q_2_CLR_100_. Q_2_ alone or with a companion drug given for 5 days prevented relapse in all footpads (Fig. 1D). The limit of detection was 1 CFU.

### Experiment 2: To determine the impact of the host immune response on the sterilizing activity of Q203-containing regimens and the selection of Q203-resistant mutants

Experiment 2 was conducted to determine if a five-day treatment with 2 mg/kg Q203 (Q_2_) daily, alone or in combination with 25 mg/kg BDQ (Q_2_BDQ_25_) daily or 12.5 mg/kg CFZ (Q_2_CFZ_12.5_) daily can successfully treat *M. ulcerans* footpad infection in SCID-beige mice without selection of Q203-resistant mutants (Table S2).

#### Footpad swelling and CFU count results up to 4 weeks post-treatment

One day after infection, the mean (± SD) CFU count was 4.17 ± 0.22 log_10_ CFU/footpad. At the start of treatment (D0), 40 days after infection, the mean footpad swelling was 2.5 (Fig. 2A), the mean CFU count was 6.26 ± 0.35 log_10_ CFU/footpad for the mice allocated to the Q203-containing regimens and 6.56 ± 0.29 log_10_ CFU/footpad for the mice allocated to the R_10_CLR_100_ control group (Fig. 2B). In all Q203-containing treatment groups, a sharp reduction in footpad swelling to a median swelling grade of 0.25 after one week of treatment was observed; in the Q_2_CFZ_12.5_ group, 19 of 34 footpads had a swelling grade of 0, and in the Q_2_ and Q_2_BDQ_25_ groups, 7 of 34 and 8 of 34 footpads, respectively, had a swelling grade of 0. In contrast, the swelling of untreated mouse footpads continued to increase such that the mice reached the humane endpoint for euthanasia at Week 1, while the control R_10_CLR_100_ regimen decreased footpad swelling after one week to a median swelling grade of only 1.5, which was still significantly higher than in the Q203-containing regimens (p<0.001). After the end of treatment in every group receiving Q203 the median swelling grade was 0, whereas the swelling started to increase in R_10_CLR_100_-treated mice again, reaching median swelling grades of 2.1 at Week 2 and ≥3 at Week 3, mandating that all R_10_CLR_100_-treated mice be euthanized per protocol at Week 4. Footpads in the Q203-treated groups maintained a swelling grade of approximately 0 until the end of the experiment. At Week 3 only 2 of the 24 footpads from mice treated with a Q203 containing regimen were culture-negative.

**Fig 2.**
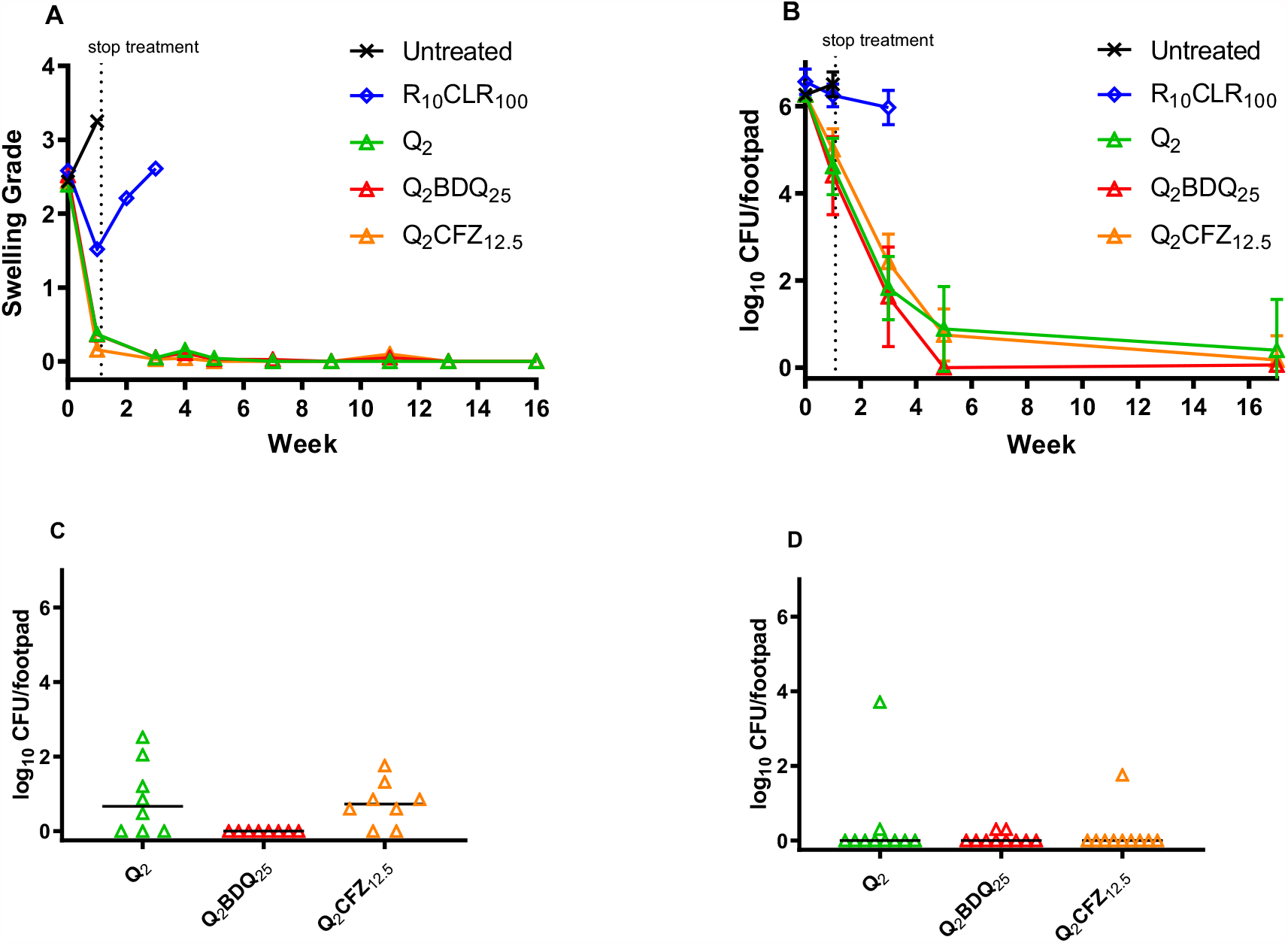
Footpad swelling grade and microbiological outcome in SCID-beige mice in response to treatment. Median swelling grade in SCID-beige mice during and after treatment with the indicated regimens and number of doses in parentheses (A). Log_10_ CFU/footpad in SCID-beige mice in the indicated regimens during the study (B). Log10 CFU/footpad five weeks after the start of treatment (C). Log10 CFU/footpad 17 weeks after the start of treatment (D). R, rifampin, CLR, clarithromycin, Q, telacebec, BDQ, bedaquiline, CFZ, clofazimine. Numbers in subscript indicate the dose in mg/kg. D0, day 0 or the beginning of treatment.

Trends in footpad CFU counts largely mirrored the swelling grades, although there was not as much discrimination between the R_10_CLR_100_ control group and the Q203-treated groups at Week 1 and the minimum values were not reached in Q203-treated groups until Week 5 (4 weeks after treatment completion) or later. Three weeks after the end of treatment, when all R_10_CLR_100_-treated mice were euthanized per protocol, their footpad CFU counts had still not decreased and were near the pre-treatment baseline. In contrast, in Q203-treated mice, footpad CFU counts continued to decline after dosing ended and, by Week 5, all footpads in the Q_2_BDQ_25_-treated group were culture-negative, in the Q_2_CFZ_12.5_ group two footpads (from two different mice) were culture-negative, and in the Q203 alone group three footpads (including both footpads in one mouse) were culture-negative (Fig. 2C). The differences between the Q_2_BDQ_25_ and the Q203 alone and Q_2_CFZ_12.5_ groups were statistically significant (p <0.03). The median footpad swelling grade as well as the mean CFU count in mice treated with any Q203-containing regimen was significantly (p<0.0001) lower than those in the R_10_CLR_100_ control group. The limit of detection was 3 CFU per footpad.

#### Relapse

Five SCID-beige mice of each Q203-containing treatment group were held for 17 weeks after starting treatment. No mouse showed a relapse in footpad swelling. In the group receiving Q203 alone, 2 out of 10 footpads had detectable CFU, in the Q_2_CFZ_12.5_ group one footpad had detectable CFU, and in the Q_2_BDQ_25_ group 2 of 10 footpads had 1 CFU each (limit of detection, 1 CFU) (Fig. 2D).

#### Resistance detection

The *M. ulcerans* isolates from the eight SCID-beige footpads with the highest CFU counts at Week 5 (4 weeks post-treatment) were selected for Q203 susceptibility testing. Two of the four footpad isolates from mice receiving Q203 alone showed resistance (40% and 2.2% of total CFU on 3 ng/ml, 40% and 3.9% of total CFU on 0.3 ng/ml, respectively) (Table S3). The Q203-containing plates from two Q_2_CFZ_12.5_-treated footpads were contaminated and were not assessable, whereas the isolates from the other two footpads from Q_2_CFZ_12.5_ -treated mice did not show resistance.

## DISCUSSION

The standard treatment for BU is an 8-week course of RIF+CLR (3), an improvement compared to the previously recommended combination of RIF+STR, as the new regimen is fully oral and avoids the previously required injections. A shorter regimen remains a high priority in the search for a better treatment of BU (19, 28). Q203, a potent inhibitor of the respiratory chain of *M. ulcerans*, is a promising candidate for ultra-short, and possibly single-dose, treatment for BU (17, 19). It has proven to be capable of significantly shortening the treatment duration in mouse footpad models (16, 17, 19, 29). Furthermore, a clinical trial showed no serious adverse effects from Q203 when taken daily for 2 weeks in a small group of TB patients (24). In January of 2021, the US Food and Drug Administration granted an orphan drug designation to Q203 for BU treatment (http://www.koreabiomed.com/news/articleView.html?idxno=10169), providing further incentives for clinical development of Q203 for this indication.

In the present study in *M. ulcerans*-infected BALB/c mice, Q203 given alone or in combination with BDQ or CFZ for 3 days was sufficient to render all footpads culture-negative 15 weeks after start of treatment. When Q203 was administered alone or in combination with BDQ, all footpads were culture-negative even five and a half weeks after the last dose. The total dose of Q203 amounted to 6 mg/kg. To our knowledge, this is the lowest total dose of Q203 to render all footpads culture-negative in a mouse footpad infection model.

No tested combination including Q203 at 2 mg/kg rendered all footpads culture-negative after a single dose. For this purpose, higher doses or other companion drugs may be necessary to achieve a single-dose cure. In BALB/c mice, none of the combinations tested was significantly more effective than Q203 alone.

To compare the sterilizing potency of Q203 and combination regimens in immunocompetent BALB/c mice and immunocompromised SCID-beige mice we analyzed the swelling grades and CFU counts in mice of each strain treated with the same regimen. All Q203-containing treatments rapidly resolved footpad swelling in SCID-beige mice, as in BALB/c mice. However, the rate of bacterial killing and the extent of bacterial eradication, while great in both strains, were not as great in SCID-beige mice. Nevertheless, by Week 15, all groups showed a relapse-free survival of at least 80% in SCID-beige mice, this result shows, that a 5-day treatment with Q203 alone (total dose of 10 mg/kg) or with a companion drug was sufficient to render the majority of footpads culture-negative and prevent relapse, even in mice lacking an adaptive immune response. At Week 5, the difference between Q_2_BDQ_25_, which rendered all footpads culture-negative, and Q203 alone, which rendered 3 of 8 footpads culture negative, and Q_2_CFZ_12.5_, which rendered 2 of 8 footpads culture negative, was statistically significant. However, at the end of the follow-up period, the difference in CFU counts was not significant.

The difference between BALB/c mice, in which all footpads were culture-negative, and SCID-beige mice, in which no group had all footpads culture-negative, is striking. This difference illustrates the importance of the immune system in the successful clearance of *M. ulcerans*. Our results suggest that treatment of BU in immunosuppressed patients, for example in the case of HIV coinfection or patients undergoing iatrogenic immunosuppression, may need to be adjusted to reach a complete cure in all patients when using Q203. The continued reduction in CFU counts supports a conclusion that the prolonged activity of Q203 is primarily due to drug effects, but the incomplete sterilizing activity of the tested regimens in SCID-beige mice illustrates the contribution of the adaptive immune system to the overall treatment response. Further research might be needed in immunodeficient mice to find a better treatment for patients, who are immune-deficient.

We did observe selection of Q203 resistance during Q203 monotherapy in SCID-beige mice. No resistance was observed when Q203 was combined with CFZ. Although only a limited number of isolates was assessed, this finding suggests that a companion drug might be beneficial to prevent emergence of Q203 resistance in very immunocompromised hosts. However, it should be recognized that SCID-beige mice represent an extreme state of immunosuppression and the majority of these mice receiving monotherapy appeared to be cured. Therefore, the pros and cons of monotherapy versus combination therapy will require careful consideration. The overall positive outcomes in SCID-beige mice, the simplicity of monotherapy and its reduced cost and potential for adverse effects, as well as the very low likelihood of human-to-human transmission of Q203-resistant BU (30, 31), should it occur, would seem to favor monotherapy. Nevertheless, further investigations may be warranted to identify an optimal companion drug that could better assure the efficacy of ultra-short (i.e., 1-3 dose) regimens across the spectrum of host immune competency (11, 32).

In summary, we identified the lowest-yet published total dose of Q203 leading to a complete healing of all affected footpads in this mouse model. Surprisingly, none of the companion drugs evaluated, contributed an additional treatment-shortening effect in BALB/c mice, nor a higher overall rate of cure in SCID-beige mice. Our studies once again show the treatment-shortening capability of Q203 but also raise the question of the optimal treatment for immunosuppressed individuals with BU.

## Materials and Methods

### Bacterial strain

For this study *M. ulcerans* strain 1059, which was originally isolated from a patient in Ghana, was used (33).

### Antibiotics

Q203 and BDQ were kindly provided by the TB Alliance. CFZ and RIF were purchased from Sigma-Aldrich (St. Louis, MO, USA). CLR was purchased from the Johns Hopkins Hospital Pharmacy. Q203 was prepared in D-α tocopheryl polyethylene glycol 1000 succinate solution. BDQ was formulated in acidified 20% hydroxypropyl-β-cyclodextrin solution. CFZ, RIF and CLR were prepared in a sterile 0.05% (wt/vol) agarose solution in distilled water.

### Mouse infection

Female 6-week-old Fox Chase SCID-beige and female BALB/c mice (Charles River Laboratories) were inoculated subcutaneously in both hind footpads with 0.03 ml of a culture suspension containing *M. ulcerans* 1059. Treatment started 5-to-6 weeks (D0) after infection when the mice had footpad swelling of grade 2-3 on a scale of 0 to 4 (27).The CFU counts at implantation were evaluated on six footpads from three mice on the day after infection (D-40 and D-34) and at the start of treatment (D0) to determine the infectious dose and the pretreatment CFU counts, respectively. The response to treatment was determined by plating each hind footpad from mice at predetermined time points.

### Treatment

Mice were treated 5 days per week with 0.2 ml per dose by gavage. Control mice treated with RIF and CLR were treated for 10 days continuously without interruption. Drug doses were chosen based on mean plasma exposures (similar area under the curve [AUC] in 24h after dose) similar to human exposures at standard doses (16, 17). All animal procedures were conducted in adherence with the Animal Welfare Act and Public Health Service Policy and other national and international guidelines. The procedures were approved by the Johns Hopkins University Animal Care and Use Committee.

#### Experiment 1

BALB/c mice (n=191 mice) were infected 5 weeks before the treatment start. The control regimen was given in the form of RIF_10_CLR_100_ for 10 consecutive days. One group of mice (6 mice) received no treatment, 6 footpads were collected one week after the beginning of treatment.

To each study group (i.e. Q203-containing groups), 14 BALB/c mice were allocated. Three groups received 2 mg/kg Q203 alone, three groups received 2 mg/kg Q203 plus 25 mg/kg BDQ, three groups received 2 mg/kg Q203 plus 12.5 mg/kg CFZ, three groups received 2 mg/kg Q203 plus 100 mg/kg CLR. Each treatment was given once daily in one group for 5 consecutive days, in one group for 3 consecutive days and in one group for one day. Six footpads per group were harvested for determination of CFU counts at 1 week and 6 weeks after the start of treatment; and 16 footpads per group were harvested 15 weeks from the start of treatment.

#### Experiment 2

SCID-beige mice (n=76 mice) were infected 6 - 7 weeks before the treatment start. The control regimen was given in the form of R_10_CLR_100_ treatment for 5 consecutive days. Three mice were held for another week without treatment. During this time the swelling increased and reached the threshold for euthanasia per protocol. At this point the mice were sacrificed and the footpads harvested for CFU counts.

To each study group (i.e., Q203-containing groups), 17 SCID-beige mice were allocated. Each group was treated once daily for 5 consecutive days. One group received 2 mg/kg Q203 alone, one group received 2 mg/kg Q203 plus 25 mg/kg BDQ, one treatment group received 2 mg/kg Q203 plus 12.5 mg/kg CFZ. At 1, 3 and 5 weeks after the start of treatment, 8 footpads per group were evaluated for response to treatment. Five additional mice from each treatment group were held for 17 weeks from the start of treatment before each footpad was harvested.

### Evaluation of treatment response

Treatment response was evaluated by (i) change in footpad swelling, as previously described (27), and (ii) change in log_10_-transformed CFU counts. The inflammatory swelling was evaluated weekly and scored as a swelling grade between 0 and 4, where a swelling grade of 0 corresponds to a normal footpad, swelling grade 1 to a swollen but non-inflammatory footpad, swelling grade 2 to inflammatory swelling of the footpad, swelling grade 3 to swelling of the entire hind foot and swelling grade 4 to an inflammatory leg swelling and ulcerations.

To determine footpad CFU counts, the footpad was thoroughly disinfected with 70% alcohol swabs and afterwards the footpad tissue was removed. The tissue was then mechanically finely minced and suspended in 1.5 ml sterile phosphate-buffered saline (PBS). Undiluted and serial 10-fold diluted aliquots (each 0.5ml) were cultured on Middlebrook 7H11selective agar, supplemented with 10% oleic acid-albumin-dextrose-catalase (OADC), at 32°C for up to 10 weeks before CFU counts were calculated.

Relapse was evaluated by holding mice for 15 weeks (Experiment 1) or 17 weeks (Experiment 2) from the start of treatment. After the end of treatment, footpads were examined every 2 weeks until the final time point was reached. Mice that re-developed footpad swelling with a lesion index ≥3 were sacrificed and the footpads plated for CFU counts. For mice reaching the final endpoint, the entire footpad homogenate was plated.

### Resistance testing in SCID-beige mice

To evaluate for selection of Q203-resistant mutants in Experiment 2, isolates were collected from culture plates from each of four mice treated with Q203 alone or Q_2_CFZ_12.5_ at the Week 5 time point. Each isolate consisted of colonies from one plate that were pooled, homogenized and then re-plated onto drug-free 7H11 agar and agar containing Q203 concentrations of 0.3 ng/ml and 3 ng/ml (4 times and 40 times MIC, respectively). The isolate was determined to be Q203-resistant if the proportion obtained by dividing the CFU count on Q203-containing plate by the CFU count on drug-free plates was ≥ 0.01.

### Statistical analysis

GraphPad Prism 9 was used to compare CFU counts and swelling grades at week 1 in Q203-treatment groups to the R_10_CLR_100_ and untreated control groups using one-way analysis of variance with Dunnett’s posttest to adjust for multiple comparisons. For the following timepoints we used one-way analysis of variance (Kruskal-Wallis test) corrected for multiple comparisons using Dunn’s test.

## Acknowledgments

This study was supported by the National Institutes of Health (R01-AI113266). O.K. was supported by a personal grant from the Bernhard-Nocht-Institute for Tropical Medicine, Hamburg, Germany. We gratefully thank the TB Alliance for providing Q203 and bedaquiline.

